# Dynamic Predictive Spatial Encoding of Motor Intentions In Area V6A of the Posterior Parietal Cortex

**DOI:** 10.1101/2025.02.06.636800

**Authors:** Antonio Roberto Buonfiglio, Stefano Diomedi, Matteo Filippini, Patrizia Fattori, Ivilin Peev Stoianov

## Abstract

To compute motor plans or intentions, the nervous system must translate target locations into body-centered coordinates. Visual stimuli, however, are sensed in retinotopic coordinates, which shift with eye movements. Furthermore, sensorimotor delays necessitate predictive processing. How does the brain compute timely gaze-invariant target locations? The dorsal visual pathway encodes spatial intentions, yet the underlying dynamic mechanisms remain elusive. Using multilevel analysis, we characterized intention coding in area V6A of the Posterior Parietal Cortex during delayed reaching tasks under diverse gaze-target conditions. We revealed a consistent population-level intention coding in V6A as eye positions changed. Next, we identified differential single-cell encoding of gaze and reaching targets in retinotopic, gaze-posture, and body-centered coordinates and elucidated the dynamical spatial normalization. Finally, we demonstrated context-dependent predictive spatial encoding in V6A, advancing our understanding of the temporal evolution of predictive visuomotor transformations during motor planning.

## 1 Introduction

Successful interactions with the environment requires the nervous system to determine object locations relative to the body [1–3]. However, visual stimuli are initially encoded in retinotopic coordinates, which shift with each eye movement. Transforming these representations into a stable body-centered reference frame is a fundamental component in sensorimotor integration [4–6] and spatial cognition [7, 8]. Understanding these transformations and their dynamics is critical for clinical applications, brain-machine interfaces, and neuroprosthetics [9–12].

The Posterior Parietal Cortex (PPC), situated at the apex of the dorsal visual stream, plays a key role in encoding spatial intentions and performing visuomotor transformations [13, 14]. Early evidence demonstrated that primate area 7 encodes behaviorally relevant locations [15]. Imaging and clinical data confirm the PPC’s role in spatial cognition and motor control in the human brain [16, 17]. By integrating visual, somatosensory, and motor signals, the PPC computes spatial goals and dynamically updates motor plans for goal-directed actions [2, 18, 19]. Although previous research has identified intention coding in the PPC, particularly in the Parietal Reach Region, area *V* 6*A* [20], and area 7 [21], the computational mechanisms underlying spatial transformations for intention encoding remain poorly understood. One influential proposal, gain-field modulation, suggests that neurons encoding locations in retinotopic coordinates scale their response with gaze position. An alternative, simpler mechanism assumes that retinotopic coordinates and gaze position are linearly combined to obtain body-centered representations [1, 13].

Predictive coding has emerged as a fundamental principle of brain function [22]. This framework posits that the brain maintains hierarchical internal models to generate predictions about current and future events, down to the sensory level. By doing so, predictive coding facilitates efficient processing and timely interactions with dynamic environments [23, 24]. In motor control, this theory explains how the brain anticipates sensory consequences of actions, compensates for inherent sensorimotor delays [25], and ensures smooth, coordinated movement [26, 27]. However, despite its theoretical appeal, direct neural evidence supporting predictive coding remains limited, particularly with regard to context-predictable stimuli in motor planning.

Here, we investigate how area V6A, located on the anterior bank of the parieto-occipital sulcus (**Fig**.1a), encodes and transforms spatial intentions during the preparatory period of delayed reaching movements under various fixation and target conditions performed blockwise allowing us to investigate both spatial transformation and predictive coding. Specifically, we test whether V6A neurons consistently encode target locations in a body-centered reference frame, even when targets appear in different retinotopic and gaze-posture locations. Furthermore, we examine whether V6A activity supports predictive coding by anticipating target locations before cues become available, challenging traditional stimulus-driven models of motor planning. We applied a multilevel approach to analyze neural data recorded from monkeys performing a delayed reaching task under three different fixation conditions [28]. First, we assessed the consistency of intention coding at the population level using multiple deep neural network decoders trained on neural activity from specific conditions and trial periods, then tested their generalization across all conditions and periods. Next, we conducted single-cell analyses to characterize response dynamics, and identify distinct subpopulations encoding target locations in retinotopic, gaze-posture, and crucially, body-centered reference frames.

**Fig. 1.**
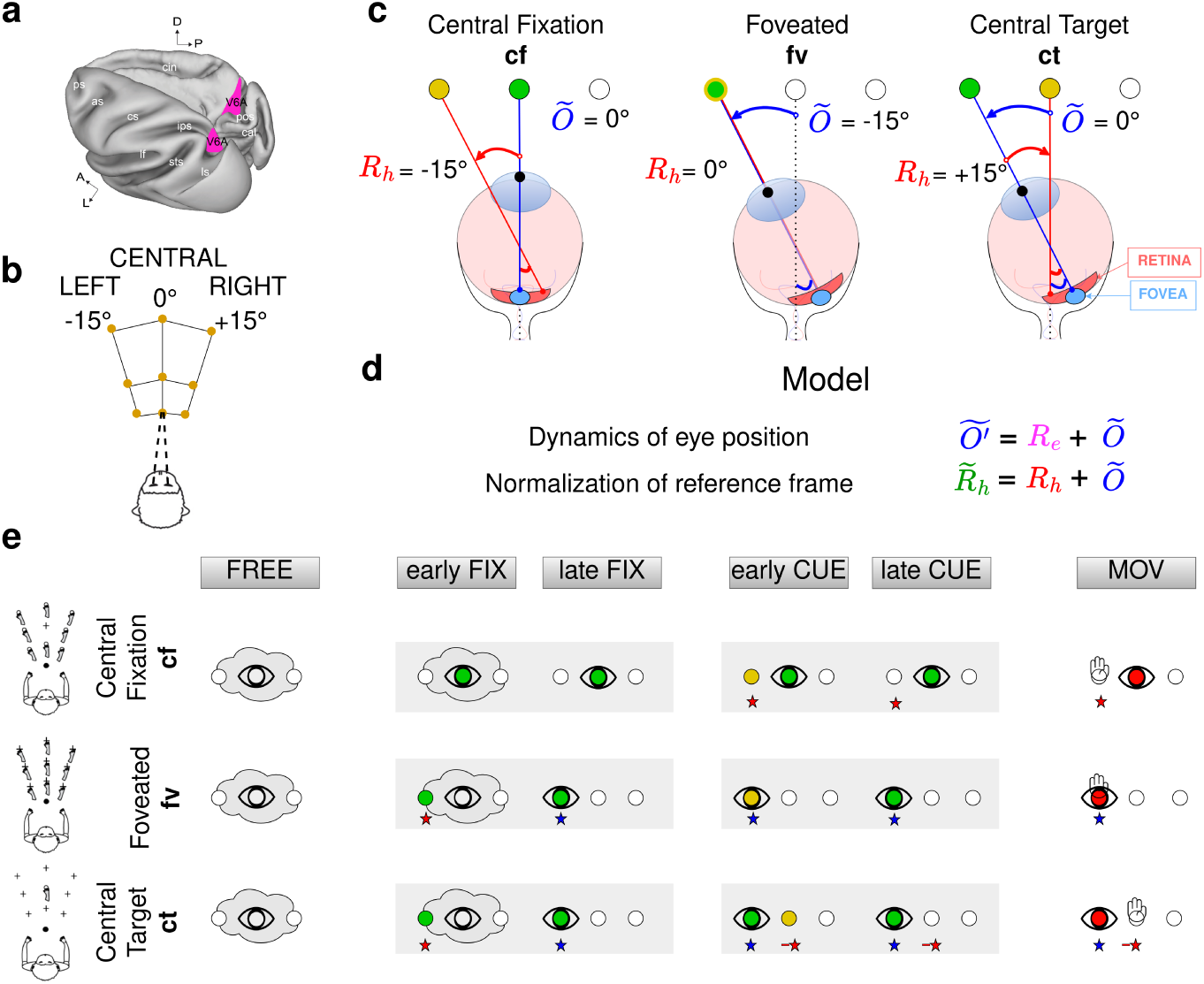
Disentangling retinotopic, gaze, and body-centered coordinates. **a**. Dorsal view of the left hemisphere and medial view of the right hemisphere of the macaque brain, highlighting area V6A. **b**. Reaching movements were performed in darkness toward one of nine possible targets located at eye level in front of a head-and-torso retrained monkey [28]. **c**. Task conditions:*Central fixation*: The eyes fixate a central location (green dot), meaning the retinotopic coordinates of the reaching target *Rh* coincide with its normalized body-centered coordinates (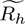, yellow dot). *Foveated targets*: Reaching targets are positioned at the origin of the retinotopic reference frame (the fovea), so their body-centered coordinates coincide with gaze posture *_O_*^-^. *Central target* : Reaching targets are always located at the origin of the body-centered reference frame, meaning their retinotopic locations correspond to the mirrored gaze posture. **d**. Computation of the predicted post-saccade gaze posture Õ*′* based on the retinotopic gaze-target *Re*, and computation of the body-centered coordinates of reaching-targets. **e**. Trial evolution across the three condition, four task phases, and two phase periods. *Abbreviations*: as, arcuate sulcus; cal, calcarine sulcus; cs, central sulcus; cin, cingulate sulcus; ips, intraparietal sulcus; lf, lateral fissure; ls, lunate sulcus; pos, parieto-occipital sulcus; ps, principal sulcus; sts, superior temporal sulcus; D, dorsal; P, posterior; A, anterior; L, lateral.

Our findings reveal that V6A acts as a computational hub, integrating retinotopic and gaze posture signals, performing spatial transformations, and maintaining gaze-invariant representations for motor control. Crucially, we provide direct neural evidence for predictive spatial coding in V6A: neurons encode target location well before sensory cues appear, demonstrating an anticipatory mechanism shaped by task context. These findings not only advance our understanding of visuomotor transformations in the PPC but also have practical implications for neural interfaces. Decoding V6A activity could enable brain-machine interfaces to extract body-centered target representation even before actual event occurrence, eliminating the need for gaze-tracking devices and enhancing real-world neuroprosthetic applications.

## 2 Results

### 2.1 Computational framework and experimental setup for motor intention encoding

We build on the notion that visuospatial information in V6A is encoded within a unified binocular reference frame that integrates signals from both eyes [29, 30]. We hypothesize that V6A encodes motor intentions across two complementary spatial domains–extrinsic and intrinsic–both represented in angular metrics with horizontal and vertical components. Without loss of generality, we focus here on the horizontal component. In the extrinsic (sensory-level) domain, visual stimuli are encoded in binocular retinotopic coordinates *R*, including both gaze and reach targets (*R_e_* and *R_h_*, respectively). In the intrinsic (motor-level) domain, spatial information is represented in body-centered angular coordinates (denoted by the symbol ∼), facilitating the kinematic computations required for motor planning [3, 14]. This includes gaze posture *Õ*, assuming a stable body-head relationship in our setup and reach targets transformed from retinotopic to head-centered coordinates 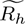. At the neural level, we assume that all these spatial domains are encoded using monotonic rate coding. Note that place coding–that underlies the gain-field modulation mechanism–can be easily converted to rate coding [31]. Rate coding is more efficient, easy to transform, and used to encode motor signals [31].

We then propose that V6A performs two key computations: reference frame transformation and predicting future gaze posture. Under the assumption of monotonic rate coding of both angular domains, retinotopic reach targets are converted into body-centered coordinates using gaze posture as a modulatory signal as follows: *R̃_h_* = *R_h_*+*Õ* (**Fig**.1d). In turn, retinotopic gaze-target coding (*R_e_*) enables direct prediction of post-saccadic gaze locations, *Õ*′ = *Õ* + *R_e_* [32, 33].

By linking the extrinsic and intrinsic domains, our framework provides an efficient mechanistic neural-level account of how V6A integrates sensory input and oculomotor signals to construct stable, predictive motor plans.

To disentangle the neural representation of retinotopic coding, gaze posture, and body-centered coordinates during reach preparation, we analyzed single-cell recordings from area V6A in monkeys performing delayed reaching tasks under various conditions (**Fig**.1b,c). Each trial began with a 1000 ms free-viewing period in darkness, during which the monkey held a start button located in front of its torso. This was followed by the onset of a fixation cue (green LED). After 1000 ms, the reach target was briefly illuminated (150ms, yellow), followed by a delay period (either 1000 or 1500 ms, randomly chosen). A reaching movement was then triggered by a change in the fixation cue color (green to red). To probe spatial representations, we designed three task conditions: (1) **central fixation** (*cf*), where retinotopic targets varied and were dissociated from central gaze; (2) **foveated** (*fv*), where retinotopic targets varied but coincided with gaze; (3) **central target** (*ct*), where fixation varied, but the reaching target remained constant. Key task events segmented each trial into four phases, corresponding to major task stages. Neural activity was extracted during two 150 ms periods per phase–one early, one late–to capture spatial coding dynamics before and after saccades, as well as early and late after reach cue onset. The spatiotemporal characteristics of fixation cues, gaze posture, and reaching target are illustrated in **Fig**.1e and further detailed in Methods.

To investigate predictive spatial encoding, we designed the experimental conditions in structured blocks, allowing subjects to infer upcoming events. This structure enabled us to assess how V6A neurons anticipate spatial transformations by encoding future gaze posture and reach targets before predictable spatial information becomes available. Specifically, we examined whether neural activity during the fixation period reflects an internal prediction of the post-saccadic gaze position and body-centered reach target location, providing evidence for predictive coding mechanisms in V6A.

### 2.2 Population-level spatial coding

We first examined how the V6A neuronal population encodes spatial domains across conditions and trial phases, assessing whether it employs a homogeneous code. To do so, we used a black-box approach with multiple deep networks (**Fig**.2a), training each network to decode spatial information relevant to a given condition, phase, and period (**Fig**.2b) while testing them on all data. For instance, *R_h_* varied in the early cue period of condition *cf*, whereas *Õ* varied from fixation onset in condition *ct*. In some periods, multiple spatial coordinates co-varied, as in *ct* (**Fig**.2c), whereas in others, no spatial dimension varied (e.g., during free viewing). The results, shown in **Fig**.2d (left matrix), display decoding accuracy on unseen data (Methods).

**Fig. 2.**
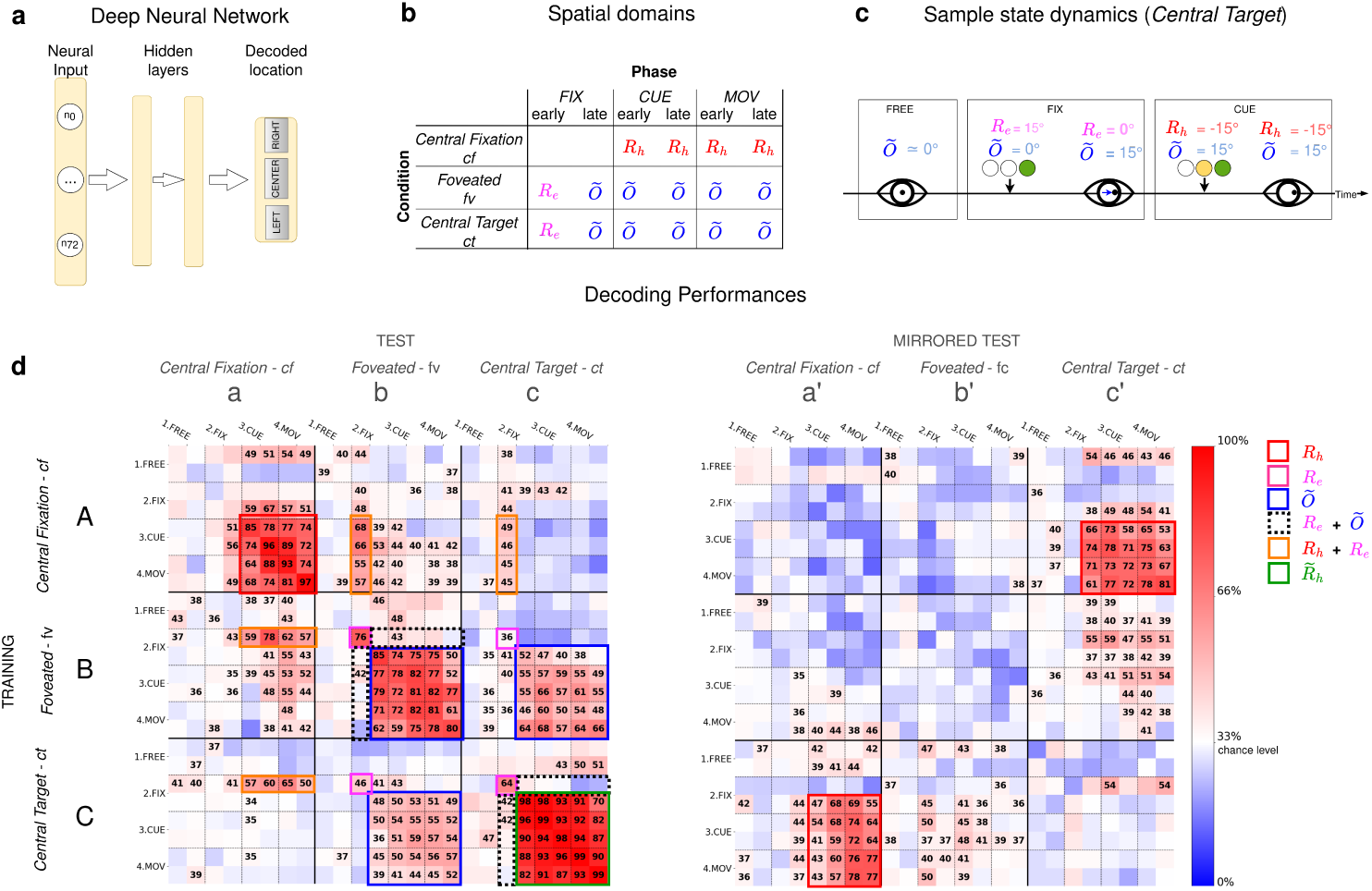
Analysis of population-level spatial coding. **a** Spatial coding at the population level was analyzed using deep neural networks trained on time-averaged activity (150 ms bins) of 73 neurons (animal A, and 50 neurons for animal B; **Suppl.Mater.**) The networks were trained to predict task-relevant spatial information on a three-level scale (Methods). **b** Spatial coordinates used to train the decoders across the different conditions, phases, and periods. **c** Evolution of spatial information in the *ct* condition. **d** Decoding performance in the normal and mirrored tests (left and right matrices, respectively). Rows indicate the condition, phase, and period used for decoder training (3 conditions x 4 phases x 2 periods, 24 decoders in total) while columns indicate the condition, phase, and period used for testing (24 tests per decoder). Cell color represents the average decoding accuracy across 10 repetitions (red to blue scale; see colorbar) with percentages displayed for values significantly different above the 33% chance level (white) after FDR-correction. Tests associated with the same type of spatial information are grouped by color boxes: red/magenta: reach/gaze targets in retinotopic coordinates, and blue for reach target in gaze coordinates.

As expected, decoding accuracy was highest when training and testing were performed on the same period (main diagonal of the matrix) and lowest when no spatial information was available, confirming the role of V6A in spatial coding. Additionally, robust generalization within each condition (red cells in the sub-matrices) indicated that V6A neurons encoded spatial information consistently during trials with congru-ent representations. More importantly, sustained spatial coding from cue onset–e.g., in condition *cf* –provided neural evidence for intention coding, or motor planning in V6A.

Crucially, our results revealed spatial compatibility between conditions. Specifically, the retinotopic target code *R_h_* in the *cf* condition was most strongly compatible with gaze-target coding (*R_e_*) just before the saccadic gaze shift in the *fv* condition. The same held for *fv*-trained decoders tested on the *cf* condition. Additionally, results suggested a link between the retinotopic target code and gaze posture *Õ*, war-ranting further analysis in the following sections. Overall, these findings support our key hypothesis that V6A encodes motor targets in a common body-centered reference frame: since *R_h_* in the *cf* condition aligns with *Õ* in the *fv* condition, they likely contribute to representing reaching targets in a normalized body-centered code 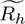= *R_h_* + *Õ* (**Fig**.1d).

The most striking result emerged from condition *ct*, where the reaching target was always central, meaning 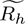= 0. Based on our framework, we predicted that neurons would encode the central reaching target through multiple spatial domains as follows (**Fig**.2c): (a) at the end of free viewing, when gaze was expected to hover around the center, the gaze target would be encoded in retinotopic coordinates *R_e_*, activated early during fixation just before the saccadic shift; (b) subsequently, as fixation stabilized, gaze posture *Õ* would corresponds to *R_e_*; and (c) finally, the central cue would activate a retinotopic reaching target code at the gaze-mirrored location, *R_h_* = −*Õ*. These components should be decodable by networks trained on the same congruent locations, such that *R_e_* = *Õ* = −*R_h_*. High decoding accuracy across relevant periods in condition *ct* would provide strong evidence for a common body-centered encodinig of reaching targets, 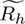= *R_h_* + *Õ*. The results in **Fig**.2d confirmed these predictions except those regarding compatibility with *R_e_*: reaching target-related signals converged on a single representation, but gaze-target *R_e_* was treated differently by V6A (see bellow).

Further supporting this interpretation, *fv*-trained decoders successfully predicted reaching target location in condition *ct*, and vice-versa. At first glance, decoders trained on *cf* and *ct* conditions seemed to reveal incompatible spatial codes, except for the retinotopic hand and gaze target codes (*R_h_*and *R_e_*). However, this matched our third prediction: the retinotopic codes *R_h_* in these conditions are opposite to each other. Indeed, the same spatial codes became compatible in a “mirror” test, where decoding was considered successful if mirrored coordinates were correctly predicted. The right matrix in **Fig**.2d, confirmed that *ct*-trained decoders successfully recognized mirrored retinotopic targets in the *cf* condition and vice versa, further supporting a common body-centered representation of reaching targets in V6A.

Finally, an unexpected result emerged: although the retinotopic encoding of gaze targets *R_e_* was strongly compatible with the retinotopic encoding of reaching targets *R_h_*, it showed only weak compatibility with the encoding of gaze posture *Õ*, which is evident in conditions *fv* and *ct* (orange boxes in **Fig**.2d). This finding suggests that V6A may process retinotopically encoded gaze and reaching targets through an overlapping subpopulation.

### 2.3 Spatial coding by single cells

While population-level analysis reveals spatial encoding in V6A, it does not clarify the role of individual neurons or whether decoding performance arises from individual neurons or their emergent combination. Additionally, training decoders on a single condition might overlook neurons with key functional roles in other conditions. This limitation is critical for clinical applications, where reliable decoding from a small subset of neurons is essential for generalization across subjects.

To address these questions, we analyzed single-cell activity. First, we assessed spatial sensitivity by comparing individual neuronal responses to left versus right stimuli across all conditions, trial phases, and periods using t-tests. The resulting contrastive discriminability patterns provided insight into each neuron’s functional role [34]. These results, visualized in **Fig**.3a, display the contrastive preference of 33 spatially sensitive neurons as a bipolar heat map, where *t*-statistics indicate left (blue) or right (red) target preferences for each condition and period (Methods).

**Fig. 3.**
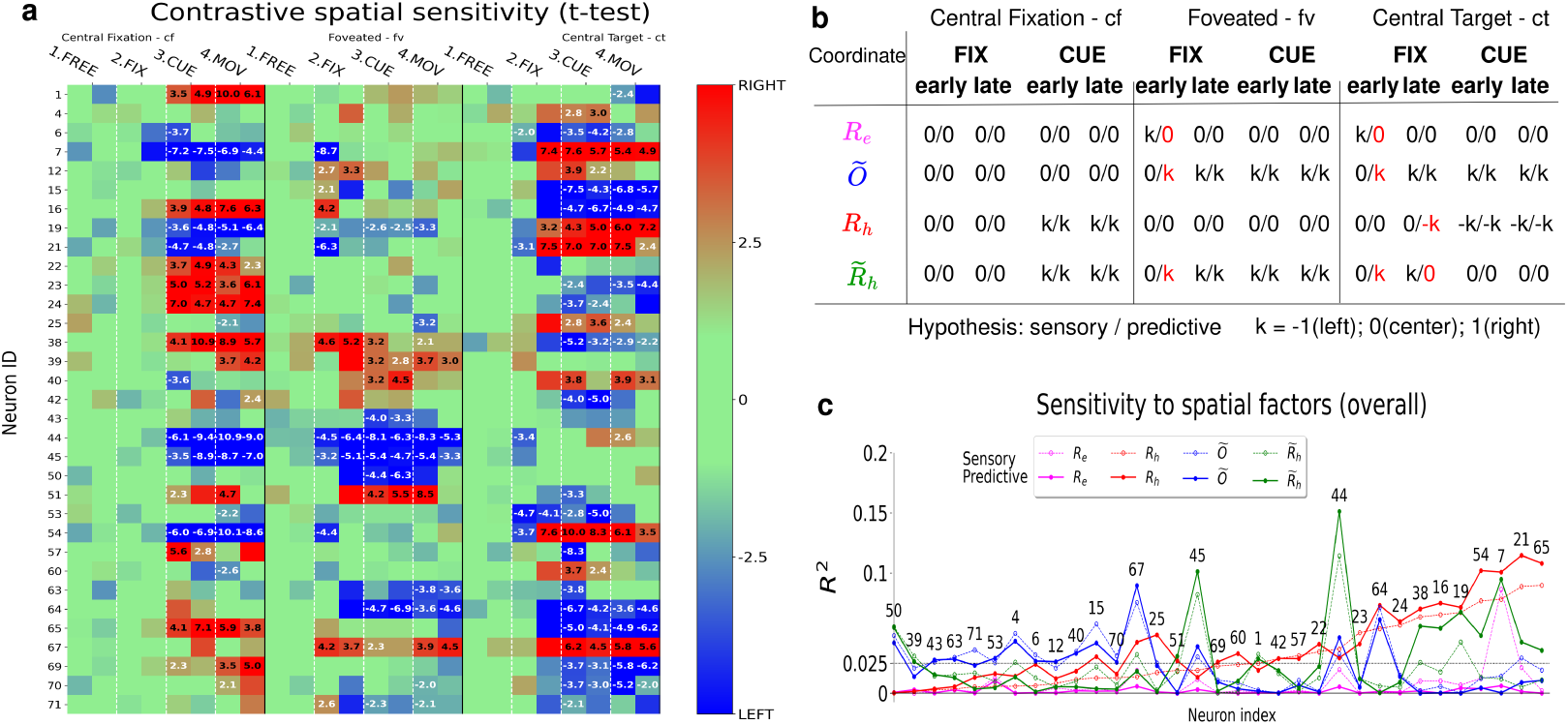
Single cell analysis. **a.** Contrastive response discriminability of 33 spatially sensitive neurons (rows) across all conditions, phases, and periods (columns), represented as *t*-values from tests comparing binned neural activity in response to left vs. right locations. Significant contrasts are also indicated numerically (see Methods and **Fig**.2b for spatial significance in each period). **b.** Spatial coordinate representations under the sensory-driven and predictive coding models. Rows correspond to spatial coordinates *Re*, Õ, *Rh*, and *R*-*h* (rows) while columns indicate condition, epoch, and period. Table cells display the coordinates under the two hypotheses. The sensory-driven model assumes a feedforward response, whereas the predictive coding model incorporates anticipatory activations when the coordinates are predictable (e.g., the reaching target position in the *ct* condition). Differences between the two models are highlighted in red. **c.** Regression fit of linear models using different spa-tial predictors: *Re* (magenta), *Rh* (red), Õ (blue), and *R*-*h* (green), under either the predictive coding hypothesis (solid lines) or the sensory-driven hypothesis (dotted lines). The horizontal black dotted line represents the threshold for identifying spatially sensitive neurons, as visualized in panel (**a**). Colors indicate the best-fitting spatial domain for each neuron: red for *Rh*, blue for Õ, and green for *R*-*h*.

The analysis highlighted several kinds of spatial sensitivity. First, **retinotopic coding** *R*: Some neurons encoded spatial location exclusively in retinotopic coordinates. For example, cell 65 discriminated left from right targets as soon as spatial cues appeared in the *cf* condition but showed no preference in the *fv* condition, except for weak discriminability during early fixation–when the fixation cue was retinotopically coded. Notably, this cell exhibited a reversed spatial preference in the *ct* condition (red in *cf* turned to blue in *ct*) indicating that in the *ct* condition it encodes the mirrored retinotopic coordinates of the reaching targets rather than gaze posture.

Second, **gaze posture encoding** *Õ*: Some neurons were selective for gaze posture rather than reaching targets. For instance, cell 50 began discriminating spatial location only after the saccade in the *fv* condition and showed no preference in *ct*, despite gaze shifting there as well–suggesting a role in gaze control rather than spatial target coding. In contrast, cell 4 exhibited contrastive activity in both the *fv* and *ct* condition, indicating that it encoded gaze posture with the purpose to represent gaze-centered reaching targets.

Third, **normalized retinotopic encoding** *R̃*: A key subset of neurons encoded targets using a normalized retinotopic code. Most notably, cell 45 exhibited contrastive spatial preference in both the *cf* and *fv* conditions. However, in the *ct* condition, its contrastive preference dropped to zero after fixation onset–likely because the mirrored retinotopic code combined with gaze posture resulted in a null, central location.

These findings highlight the diversity of spatial encoding strategies in V6A, with different neurons selectively encoding retinotopic targets, gaze posture, or a combination of both in a body-centered reference frame.

### 2.4 Predictive coding for motor planning

The contrastive single-cell analysis revealed unexpected spatial selectivity in V6A neurons. For instance, the “retinotopic” neuron 65–which was activated by rightward targets in the *cf* condition–became selectively responsive to left fixation cues in the *ct* condition. Notably, this activity emerged *before* the reaching target appeared during the cue period, effectively predicting the future presence of a centrally located cue in left retinotopic coordinates (**Fig**.3a). Such anticipatory activity was possible in our experimental paradigm, in which conditions were presented in blocks, allowing subjects to infer upcoming events. For example, monkeys could anticipate that in the *ct* condition, (i) targets would appear centrally, (ii) gaze posture would shift immediately after fixation, and (iii) the retinotopic coordinates of the next reaching target would be inverted relative to gaze posture. This predictive ability enables proactive motor planning, which is essential in real-world scenarios such as evading predators or hunting in familiar environments [35].

To systematically investigate predictive coding in V6A, we formulated two competing hypotheses regarding the representation of each spatial coordinate across different conditions, epochs, and task periods (see Methods and **Fig**.3b). These hypotheses were grounded in two contrasting theories of perception: traditional feedforward processing and predictive coding [22]. The fundamental distinction between these two theories lies in their temporal dynamics. Under a strictly stimulus-driven, feedforward framework, gaze posture would be updated only after a saccadic shift and reach target would be encoded only after the reach cue is shown. In contrast, under predictive coding, the representation of gaze posture is updated in advance [32]. We leveraged these theoretical differences to construct model-based predictions of neural activity, which we tested using linear regression analyses.

This approach identified 33 spatially sensitive cells (out of 73 analyzed in monkey A and 9 out of 50 in monkey B, **Suppl.Mater.**), where at least one model exhibited a regression fit of *R*^2^ *>* 0.025. **Fig**.3c summarizes how each spatial factor and predictability assumption influenced neural responses. Among these neurons, 48.5% were most sensitive to retinotopic target location (*R_h_*), 33.3% to gaze posture (*O_t_*), and 18.2% to body-centered target location 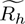. Notably, 63.6% of neurons exhibited activity best explained by predicted spatial representations, while the remaining 36.4% were better explained by current spatial coordinates.

These findings quantitatively confirm and extend insights from the contrastive analysis, demonstrating that V6A neurons encode not only sensory-driven spatial information but also internally generated predictions of spatial states. Our result provide strong evidence that predictive processing is a fundamental mechanism of sensorimotor planning, with V6A playing a central role in this anticipatory computation.

### 2.5 Functional signatures and dynamic transformations

To refine our understanding of V6A neurons’ functional roles, we examined their dynamic response patterns. A neuron’s activity was best interpreted through its pattern of model fits, which together defined a functional signature–a quantitative description of how its response aligned with different spatial coordinates. In **Fig**.4, we illustrate these functional signatures alongside the dynamic response profiles of six representative neurons.

**Fig. 4.**
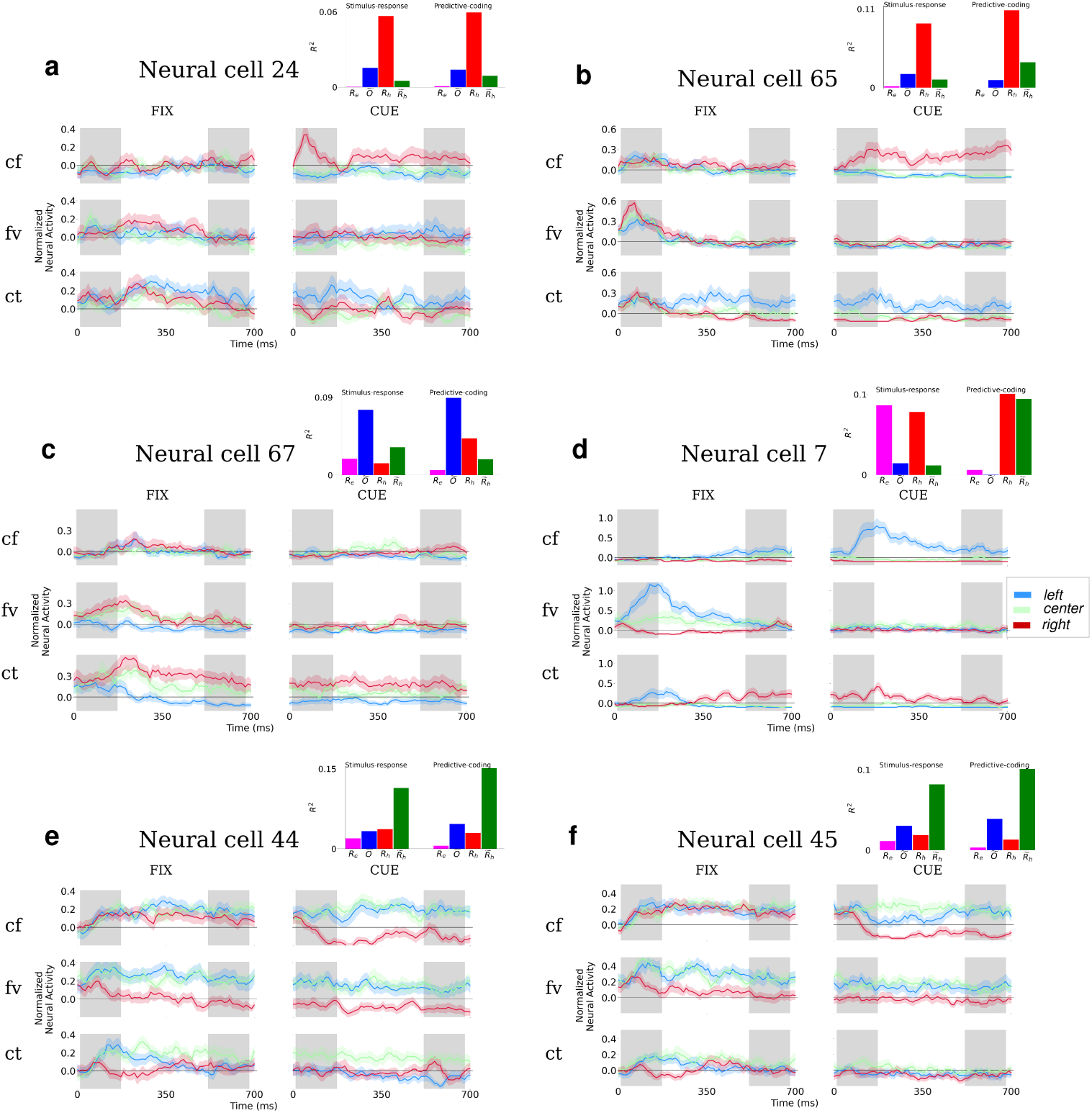
Response and signature of six cells (panels a-f). Lines visualize cell response during the fixation and cue periods under every condition, averaged over the left, central and right targets (color legend; bands show st.err.). Cell signatures include the regression fit to every spatial coordinate. Cells 24 and 65 respond to right reaching targets encoded in retinotopic reference frame. Note that under condition *ct* these correspond to left fixations. Moreover, under *ct*, these cell start responding right after the fixation cue, predictively, well before the reaching target appears. Cell 67 appears to encode gaze position under the *fv* and *ct* conditions. Cell 7 is retinotopic, encoding *Re* and later *Rh* in *ct* trials. Cells 44 and 45 compute body centered representation of the reaching targets, predictively.

Cells 24 and 65 exhibited the strongest alignment with retinotopic target coding (*R_h_*), a pattern clearly reflected in their temporal response profiles. These neurons discriminated between left and right targets during the cue phase of *cf* trials and again in the late fixation and cue phases of *ct* trials. Notably, in *ct* trials, they encoded reaching targets predictively, maintaining a retinotopic reference frame before the target was presented.

Cell 7 followed a similar but with an important distinction: it also discriminated gaze targets (*R_e_*) during the early fixation period of *fv* trials, indicating sensitivity to gaze-related spatial information. In *ct* trials, its response selectivity initially mirrored that of *cf* and *fv* trials. However, following the gaze shift, the cell reversed its response, suggesting a dynamic updating of its retinotopic encoding. This response reversal reflects a flexible transformation process, where the neuron predictively encoded the new retinotopic location of the reach target.

Cell 67 displayed a distinct pattern, being highly selective for gaze posture rather than target locations. This was evident in both *fv* and *ct* conditions, where it maintained a consistent spatial preference, encoding both the gaze target and the future gaze position. Unlike retinotopic neurons, its activity did not reverse after the gaze shift in *ct* trials, indicating a stable representation of gaze-related spatial states.

Cells 44 and 45 exhibited the most intriguing response patterns, corroborating findings from the contrastive response analysis. These neurons were highly active in *cf* and *fv* trials, showing clear spatial discrimination. In *fv* trials their selectivity emerged as early as fixation cue onset. Their strong preference for left-reaching targets in both conditions suggested they encoded body-centered target representations. However, their response was markedly different in *ct* trials. Despite discriminating between left and right cues at fixation onset, they failed to maintain selectivity during the cue period. This shift suggests that their encoding was body-centered, meaning that in *ct* trials, the target location remained centrally aligned with the body.

A closer look at their response dynamics reveals a predictive cancellation effect: Initially, these neurons responded to the fixation cue (*R_e_*), which later transitioned to encoding gaze posture (*Õ*). At the same time, they also integrated the retinotopic reaching target (*R_h_*)–but predictively, before it appeared. In *ct* trials, this predictive response had the same amplitude as gaze posture encoding but the opposite sign, leading to a net cancellation of activity.

This finding provides further evidence that predictive spatial transformations in V6A involve not just encoding expected spatial targets but also dynamically integrating multiple spatial frames to maintain a stable, body-centered reference for motor planning.

## 3 Discussion

We investigated predictive intention encoding and reference frame transformations during motor planning in area V6A, a region of the PPC at the apex of the dorsal visual stream known to play a crucial role in visuospatial processing and motor preparation [1, 13, 36]. By integrating formal analyses of reference frame transformations, population activity decoding, and single-cell analyses, we provide insights into the dynamics of motor intention encoding in this sensorimotor hub. Our findings reveal the temporal evolution of sensorimotor transformations in V6A showing that while population-level activity robustly encoded movement intentions during both motor planning and action execution, individual neurons exhibited distinct spatial representations. These neurons encoded predictively, shaped by the experimental context and task structure, gaze targets, gaze posture, and reaching targets in both retinotopic and body-centered coordinates as the gaze shifted. The dynamic spatial normalization process ensured the timely computation of gaze-invariant target positions during reach tasks across different gaze-posture conditions. These results extend foundational studies of spatial encoding and motor planning [21, 36], providing neural evidence for predictive spatial encoding in area V6A.

### Dynamic spatial encoding

V6A plays a pivotal role in visuomotor processing, particularly in motor intention planning [5, 13, 36]. The temporal dynamics of individual neurons observed in our study revealed distinct phases of intention encoding during reach task planning, transitioning from initial gaze-dependent signals to integrated body-centered motor plans, while maintaining stable intention coding at the population level. This extends prior research linking PPC activity with motor intention [21] and object-centered spatial transformations [37]. Our results corroborate and expand foundational studies that establish V6A as a hub for integrating visual and motor signals [38], reveal neuronal representations of both target location and intended movement direction [39], and provide evidence for multiple spatial representations in V6A during reach planning and execution [40]. In addition, by demonstrating the robustness of motor intention coding across diverse gaze-target conditions, we offer new insights into the neural mechanisms underlying goal-directed actions.

V6A’s ability to transform spatial information across retinotopic, postural, and body-centered reference frames is essential for coordinating complex eye and hand movements. While previous studies documented reference frame transformations in the PPC [18, 41, 42], they did not describe the temporal dynamics underlying gaze normalization. Here, we identified gaze-invariant coding at the population level and demonstrated V6A’s role in orchestrating reference frame transformations through the flexible integration of eye, head, and body posture signals. These findings extend results describing the adaptability of V6A neurons in integrating spatial features, such as gaze position, during reaching tasks [43].

### Predictive Coding

Temporal delays, inherent to neural information transmission, pose a fundamental challenge for biological motor control–a challenge the brain appears to overcome through anticipatory mechanisms that optimize the entire control process, from perceiving environmental states [22] to motor planning and action execution [3, 27, 44]. More than half of the V6A neurons analyzed in this study encoded spatial signals predictively, anticipating future target locations well before stimuli onset. These results elucidate the temporal dynamics of predictive coding in V6A, revealing how neuronal activity evolves to encode future target positions, ensuring accurate motor planning in anticipation of upcoming events.

Our findings expand previous work showing that neurons in the V6A area integrate multiple spatial reference frames during movement planning [45] that dynamically adapt their spatial encoding based on contextual demands [40, 43]. The combined contrastive analysis and dynamic profiles highlight the temporal evolution of predictive signals, suggesting that V6A actively anticipates future movement parameters. Notably, we also reveal a tight interaction between predictive coding and reference frame transformations: predictive signals in V6A were systematically coupled with predictive transitions between retinotopic and body-centered reference frames occurring in a temporally coordinated manner. Thus, predictive coding in V6A is not limited to encoding future target locations but it rather operates as a context-dependent, dynamic process that continuously maintains the appropriate spatial frame for coordinated eye and hand goal-directed movement.

The results extend predictive coding research, revealing neurons in parietal areas 7a and 5d that anticipate dynamic visual targets [25]. This aligns with predictive coding frameworks based on active inference in continuous time, which assume not only static representations of predicted and perceived stimuli but also complete perceptual dynamics encoded in the visual areas [14, 23]. In contrast, in our study, it is not visual dynamics but rather the structured design of experimental blocks and trials that enables the prediction of reach targets. Such long-term contextual information is likely provided by higher-level cortical areas such as the Prefrontal Cortex (PFC). A promising recent extension of the active inference in continuous time–hybrid active inference–offers a comprehensive explanation for these dynamic, context-dependent processes. This framework consists of a lower-level continuous sensorimotor control component, likely corresponding to PPC functions, that is dynamically coupled with higher-level discrete planning, presumably implemented in the PFC and related areas [44, 46].

### Multilevel analysis

We leveraged deep neural network decoding to provide novel insights into population-level dynamics. Integrating population decoding with single-cell investigations in a multilevel analytical framework allowed us to uncover fine-grained details of the transformations occurring in V6A and their temporal evolution, offering a more interpretable framework than conventional decoding methods. This approach also extends traditional single-cell analyses [18, 36], which, although effectively characterizing individual neural response, don’t address computational function. Conversely, decoding analysis often overlooks the contribution of single-cell activity to decoding outcomes, prioritizing decoding accuracy over functional insights [39, 47, 48].

Combining decoding and single-neuron investigations addresses the black-box limitation of artificial neural networks. Our method parallels explainability techniques for deep neural networks, aiming to formulate a cohesive functional theory explaining decoding performance across conditions. This aligns with foundational research, for example [49] that used backpropagation neural networks to simulate PPC neuronal response to retinotopic stimuli. However, while that study validated experimental observations, it did not reveal novel computational insights. Other neurocomputational approaches integrate artificial and biological data, at multiple levels, including neural and behavioral, to shed light on complex perceptual, motor, and cognitive processes [3, 35, 44, 50, 51]. Our research provides novel data for this kind of modeling effort and moreover details a specific hypothesis about the neural computations in V6A underlying the coordination of eye and hand movements during motor planning and visuomotor integration.

The integration of decoding techniques with single-cell analyses helped corroborating previous findings and revealed new details regarding the underlying neural computations. Beyond unveiling new functional capabilities of V6A, this approach provides a more comprehensive analytical framework for future research and introduces potentially more robust methods for Brain-Machine Interfaces.

### Limitations and Future Directions

Our focus was on the preparatory period, and we did not conduct a detailed analysis of sensorimotor integration during movement execution. However, the contrastive analysis suggests strong continuity in the spatial representation between planning and execution at the level of individual neurons, reinforcing the key role of persistent intention coding during action execution. Persistent goal coding may be essential for continuous sensorimotor control, supporting fine error corrections–a fundamental component in forward models [3, 23, 52]. Future research could extend these findings by analyzing neural dynamics during movement execution, particularly in relation to predictive coding mechanisms. A richer dataset–including co-registered neural activity, eye-tracking, and kinematic measurements–would enable a more granular investigation of predictive processes across task phases, potentially clarifying the role of effector-, or limb-centered reference frames [53] and help validating alternative theoretical frameworks, such as optimal control [54].

Neurocomputational models based on active inference provide a promising framework for integrating perception, planning, and control [3, 14, 23]. These models could help contextualize our findings within a broader theoretical perspective, elucidating the principles of predictive spatial coding and motor control. Expanding this approach could also facilitate the study of more complex spatial computations in ecologically valid settings, including unrestricted behaviors such as grasping [20, 46], tool use [44], and dynamic visuomotor tasks requiring continuous adaptation of action selection [55].

### In conclusion

this study provides novel insights into the neural computations underlying reference frame transformations and predictive spatial coding in area V6A. By leveraging a multilevel analytical approach, we demonstrated how V6A anticipates spatial targets in a context-dependent manner, dynamically integrating retinotopic and postural signals to optimize motor planning. These findings advance our understanding of visuomotor transformations in the PPC with relevance for neuropsychological and clinical research, and open new avenues for research on predictive motor control, computational models of sensorimotor integration, brain-machine interfaces, and neuroprosthetics.

## METHODS

### Behavioral setup

Reaching tasks were performed in darkness with the hand contralateral to the recording hemisphere. Multi-color Light Emission Diods (LED) attached on levers used to register the monkey response and located on a 2D-plane near the eye vertical level were used as fixation- and reach-targets. The cues could be presented in three horizontal isovergent locations (−15°, 0°, 15°); vergence was varied as well (17°, 11.5°, 7°) to increase spatial variability but it was not considered here.

### Neural data

We combined data obtained in separate sessions for each of the n=73 cells recorded from subject A and 50 cells from subject B. The recording sessions of each cell included 10 task repetitions for each of the nine targets under each spatial condition, for a total of 270 trials per cell. The tasks in each of the three conditions were performed in separate blocks. Within a given block, we varied only fixation and/or target location according to the setup described in Fig.1. We split the recordings in trials, extracted spike activity, and grouped the spiking activity in bins of 2 ms and z-scored the binned data for every neuron.

We restructured the neural data to build a dataset for the decoders, consisting of vectors containing the activity of all 73 cells (monkey A; 50 cells, monkey B) registered while the monkey was performing the task in all 3 conditions, 9 targets, 10 repetitions per target, 4 task phases, and two periods per phase (an early period, in the 30 − 180 ms time interval and a late period, in the 500 − 650 ms time interval post onset of of a given task phase), averaged the neural activity over each period. This resulted in a dataset comprising in total n=2160 vectors, 73-units long, each labeled with an index of the condition/phase/period/position which it represented.

### Decoders

were deep neural networks trained to predict spatial properties of a given trial, using as input vectors of neural activity. The deep networks featured two fully connected hidden layers, each containing 32 ReLU units (Fig.2a). We used supervised learning with cross-entropy loss function to train the decoders to learn to predict spatial features, which were horizontal locations over a 3-level scale: left, center, and right. We trained in total 24 different decoders (3 conditions, 4 task-phases, 2 periods per phase); the spatial features for each condition/task-phase/period are described in Fig.2b. Given the limited amount of data, decoding was assessed by applying cross-validation that included n=10 pseudo-random splits of the entire dataset into a training subset containing n=8 repetitions of each target, a validation subset containing one target repetition, and a test subset containing one target repetition, too; this resulted in 240 trained decoders in total. The trained decoders were used to make predictions for the unseen data, not only belonging to the same condition/phase/period used for training, but also for all other conditions/phases/periods.

### Spatial sensitivity

Contrastive analysis compared statistically with the help of t-tests the neural activity in the left and right trials, with the aim to assess the spatial sensitivity of each cell, in particular, the capacity to discriminate between right and left targets over time. The data for this analysis was extracted from the dataset used in the population analysis. Considering that each target was repeated 10 times and that there were three left and three right targets (one for each depth level), each t-test contrasted the neural activity of 30 left and 30 right trials. Results are shown on **Fig**.3a on a blue-to-red color scale, with green color meaning zero; t-statistics of significant t-tests are also shown as numbers (we applied FDR correction for multiple contrasts). In the figure, rows show the neural cells that strongly represent at least one spatial dimension of interest in at least one period. To identify these neurons, we conducted an additional analysis described hereafter and considered as spatially sensitive those neurons that had model fit *R*^2^ *>*= 0.025 (an arbitrary chosen threshold); see **Fig**.3c.

### Hypothesis for predictive coding

To investigate the specific functional role of individual neurons and the possibility that individual neurons respond predictively, we first schematized the sequence of events in each condition. During freeviewing, we assume that gaze hovers around a central neutral posture, *Õ* ≈ 0. Then, in the *cf* condition, the fixation cue appear at the center and after about 1000 ms, the reach target *R_h_* appears briefly at an unpredictable location. Instead, in the *fv* and *ct* conditions, the fixation cue appears first at an unpredictable location and it is encoded in retinotopic coordinates *R_e_*during the early fixation period. The perception of this target allows predicting that a few hundred of milliseconds later, during the late fixation period, gaze will be shifted to that location, i.e., *R_e_*(*t*) 1↦ *Õ*(*t* + 400*ms*) and furthermore, specifically to the *ct* condition, a reach-target will appear in the center, at retinotopic coordinates that mirror the future gaze position, i.e., *R_e_*(*t*) 1↦ *Õ*(*t* + 400*ms*) 1↦ −*R_h_*(*t* + 1000*ms*). Instead, in the *fv* condition, the reach target is flashed at the location of the fixation cue. We used the task structures to hypothesize how each spatial coordinate might drive the individual cell activity in every condition under the two predictability hypotheses, which allowed us to verify whether V6A neurons exploit the predictable task structure to prepare reach plans in advance. The hypotheses are shown in **Fig**.3b.

### Modeling single cell response

We modeled the response of each neuron using linear regressions with predictors corresponding to the different hypotheses about encoded spatial coordinate and predictability hypothesis. These models offer a more nuanced analysis than the contrastive analysis, highlighting to what extent the individual neurons encode the different spatial coordinates. More importantly, they allowed us to verify whether contextually anticipated spatial coordinates helps to explain better individual neuronal activity.

The following equations describes the linear regressions computed for each cell *c* and spatial predictor. In Eq. 1 *y̅* is the averaged neural activity over the 150*ms* time period (dependent variable); *β*_0_ and *β*_1_ are the intercept and the slope of the linear regression and *x* is target coordinate (−1 left, 0 center, +1 right). The *x*-values of all coordinate systems are described in the table shown in **Fig**.3b.

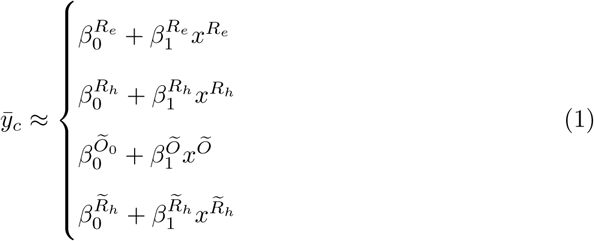

The quality of the regression models was quantified using the regression fits *R*^2^, shown in **Fig**.3c, which allowed us to determine which predictor explains best the overall response of every individual neuron under the two predictability hypotheses.

## Acknowledgments

This research received funding from the European Union’s Horizon H2020-EIC-FETPROACT-2019 Programme for Research and Innovation under Grant Agreement 951910 (MAIA) to I.P.S. and P.F. It was also supported by #NEXTGENERATIONEU (NGEU) and funded by the Italian Ministry of University and Research (MUR), National Recovery and Resilience Plan (NRRP), project MNESYS (PE0000006) – A Multiscale integrated approach to the study of the nervous system in health and disease (DN. 1553 11.10.2022). The funders had no role in study design, data collection and analysis, decision to publish, or preparation of the manuscript.

## 4 Data availability

Data, code, and further information regarding the statistical analyses are available at https://github.com/stoianov/v6a predictive spatialencoding

## Notes

https://github.com/stoianov/v6a_predictive_spatialencoding

### Competing Interest Statement

The authors have declared no competing interest.

