## Supplementary Figures for "Dynamic Predictive Spatial Encoding of Motor Intentions In Area V6A of the Posterior Parietal Cortex"

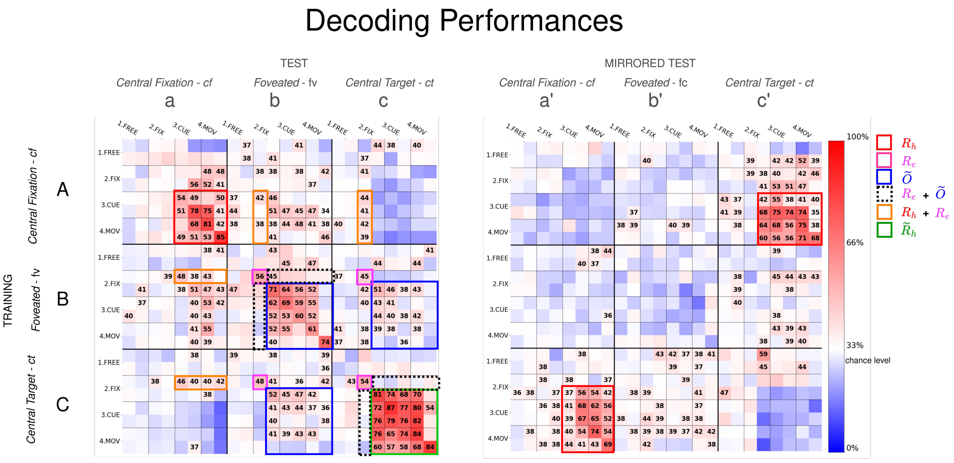


**Fig. 1** Monkey B population decoder. Decoding performance in the normal and mirrored tests (left and right matrices, respectively). Rows indicate the condition, phase, and period used for decoder training (3 conditions x 4 phases x 2 periods, 24 decoders in total) while columns indicate the condition, phase, and period used for testing (24 tests per decoder). Cell color represents the average decoding accuracy across 10 repetitions (red to blue scale; see colorbar) with percentages displayed for values significantly different above the 33% chance level (white) after FDR-correction. Tests associated with the same type of spatial information are grouped by color boxes: red/magenta: reach/gaze targets in retinotopic coordinates, and blue for reach target in gaze coordinates.

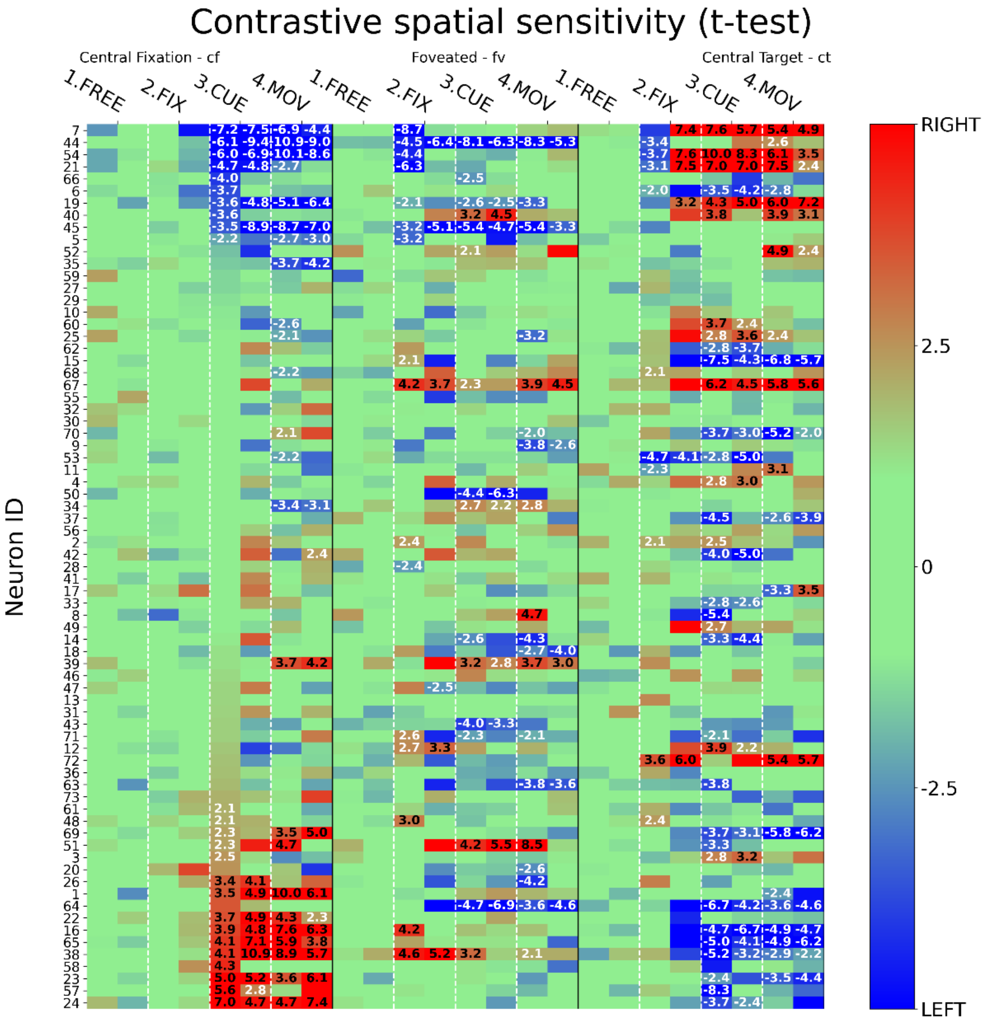


**Fig. 2** Single cell analysis of the entire neural population of Monkey A. Contrastive response discriminability of neurons (rows) across all conditions, phases, and periods (columns), represented as t-values shown on a color scale (colorbar) of tests comparing 150-ms bin neural activity in response to left vs. right locations. Significant contrasts are shown numerically.

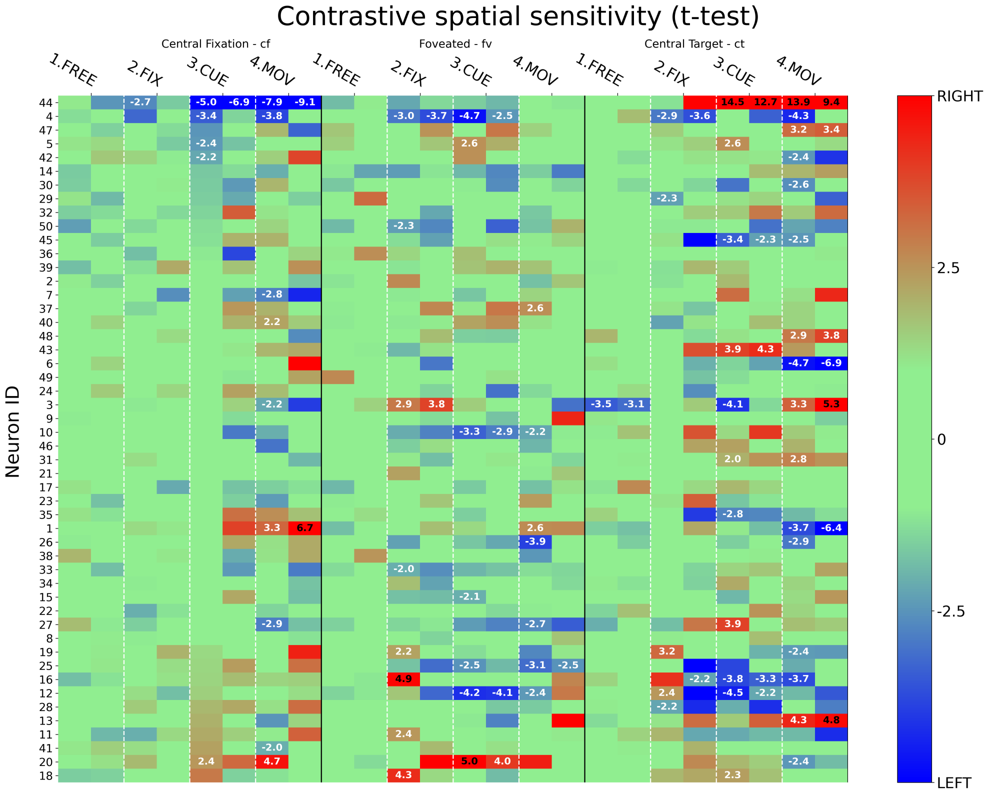


**Fig. 3** Single cell analysis of the entire neural population of Monkey B. Contrastive response discriminability of neurons (rows) across all conditions, phases, and periods (columns), represented as t-values shown on a color scale (colorbar) of tests comparing 150-ms bin neural activity in response to left vs. right locations. Significant contrasts are shown numerically.

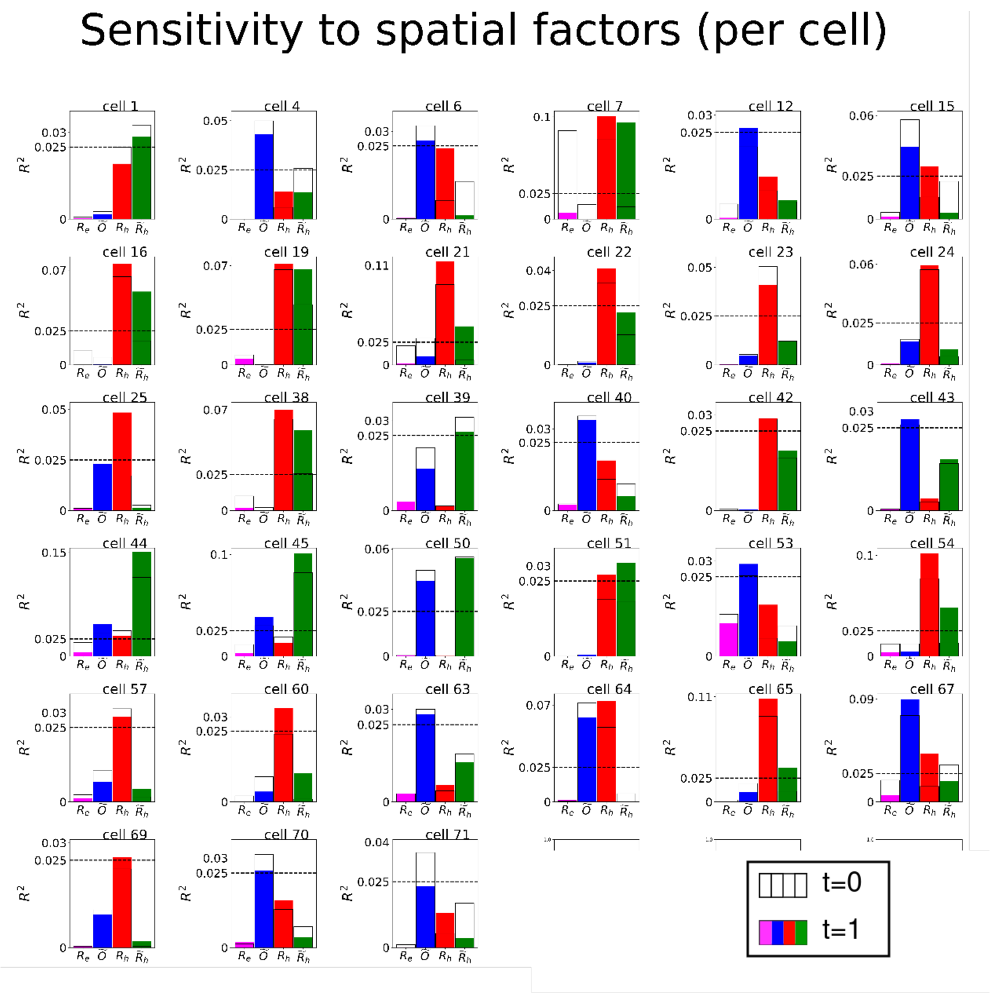


**Fig. 4** Signatures of the entire neural population of Monkey A. The signatures include the regression fit of every spatial coordinate.The horizontal black dotted line represents the threshold used for identifying spatially sensitive neurons. Colors indicate the best-fitting spatial domain for each neuron: red for $R_h$, blue for $\widetilde{O}$, and green for $\widetilde{R}_h$.

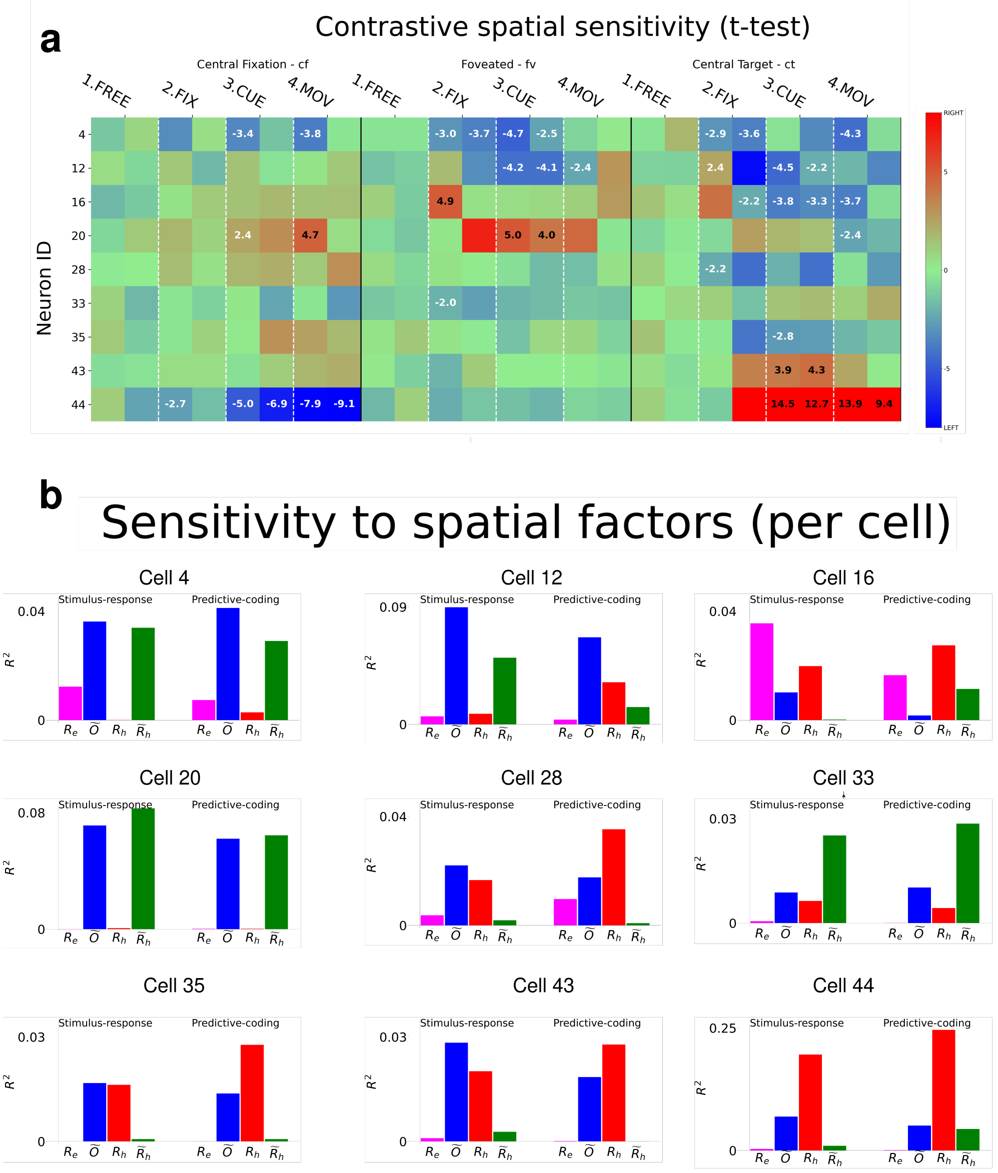


**Fig. 5** Neurons encoding spatial properties, monkey B. **a** Contrastive response discriminability of 9 spatially sensitive neurons (rows) across all conditions, phases, and periods (columns), represented as t-values from tests comparing binned neural activity in response to left vs. right locations. Significant contrasts are also indicated numerically. **b** Signatures of the neurons shows in pannel a . Cell signatures include the regression fit to every spatial coordinate.

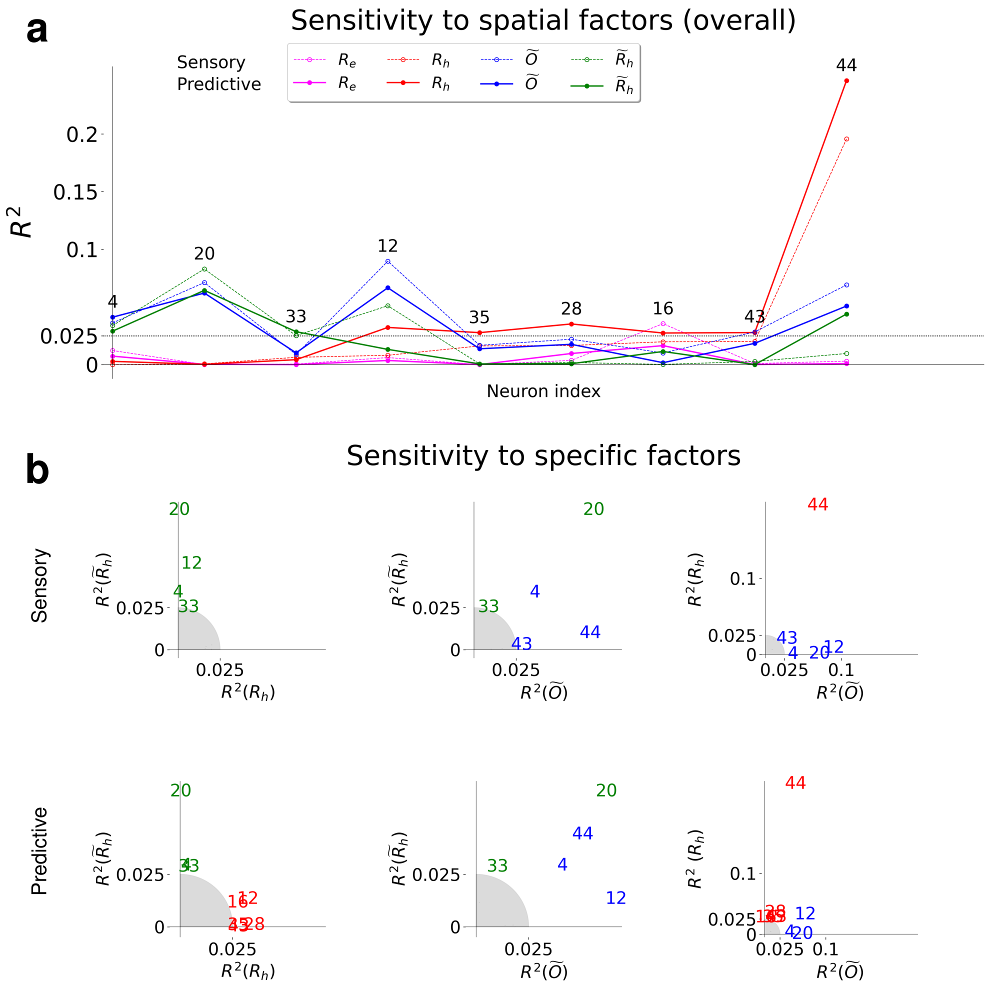


**Fig. 6** Regression analysis, Monkey B. **a** Regression fit of linear models using different spatial predictors: $R_e$ (magenta), $R_h$ (red), $\widetilde{O}$ (blue), and $\widetilde{R}_h$ (green), under either the predictive coding hypothesis (solid lines) or the sensory-driven hypothesis (dotted lines). The horizontal black dotted line represents the threshold used for identifying spatially sensitive neurons, as visualized in panel. **b** Pairwise comparisons between specific regression models. The first and second rows compare models under either Sensory-response or predictive coding hypothesis.
